# Infinite horizon control captures modulation of movement duration in reaching movements

**DOI:** 10.1101/2025.10.22.683958

**Authors:** Antoine De Comite, Hari Teja Kalidindi, J. Andrew Pruszynski, Frédéric Crevecoeur

## Abstract

Movement duration, a fundamental aspect of motor control, is often viewed as a pre-programmed parameter requiring dedicated selection mechanisms. An alternative view posits that movement duration emerges from the control policy itself. Here, we demonstrate, using infinite horizon optimal feedback control (IHOFC) and nonlinear limb dynamics that this alternative hypothesis successfully captures diverse aspects of human reaching behavior, including tradeoffs between movement duration and task parameters. Specifically, we reproduced the modulation of movement duration with varying reach distances and accuracy (Fitts’ law), and extended the infinite horizon framework to include the effect of rewards and biomechanical costs. Furthermore, our model also featured a temporal evolution of feedback responses to perturbations that resembles experimental observations, and naturally accounted for motor decisions observed when participants select one among multiple goals in dynamic environments. Together, these developments show that in many cases, movement duration may not need to be specified a priori, but instead could result from task-dependent control policies. This framework validates a candidate explanation for varied movement durations, which invites to reconsider the nature and strength of evidence for the finite horizon formulation.

## Introduction

Human movements are highly variable, and their properties depend on the context in which they are performed. While the covariation of movement duration with various task parameters such as distance to target, accuracy requirements, or rewards has been thoroughly described [1], [2], little is known about the principles by which the brain may select movement duration. Two main modeling approaches have emerged to explain this covariation of movement duration with task parameters. The first one assumes that movements are executed and controlled over a fixed duration, determined a priori, which corresponds to a finite horizon control framework [3], [4], [5]. Alternatively, a second approach proposes that movement duration emerges from the task specifications, which corresponds to an infinite horizon control framework [6], [7]. Despite the flexibility of the latter approach, its usage has remained constrained to explaining a subset of movement features such as Fitts’ law [7] or the response to visual perturbations during reaching [6]. Here, we investigated to what extent this approach was generalizable to other tasks by reproducing a larger number of experiments known to evoke changes in movement duration and by considering nonlinear arm dynamics.

Humans reaching movements are characterized by tradeoffs between movement duration and task parameters that the infinite horizon control framework could capture. A first tradeoff, described by Fitts’ law [2], relates movement duration to movement amplitude and movement accuracy. While finite horizon models were able to capture this tradeoff by predetermining a movement time (possibly optimal) [3], [4], infinite horizon controllers captured it as an emerging property of their optimization process [7], [8]. However, these infinite horizon models relied on linear plant dynamic and determined the movement end based on a threshold on the hand variance [6], [7], which differed from the experimental settings where the limb dynamics are nonlinear and movement end is often determined based on position and velocity thresholds. More recently, a second prevalent tradeoff between movement duration and reward was documented in human movements [1], [9], [10], [11]: movements are faster in presence of larger reward. This tradeoff was explained using the notion of cost-of-time [12], built on the assumption that the goal of movements is to aim for a rewarding state whose value depletes with time. The cost-of-time penalizes slow movements, while motor cost penalizes fast movements, resulting in a tradeoff that defines the optimal movement duration. Movement duration can therefore be derived from the task specification [12], [13], [14] and used as an input to finite horizon controllers [15]. In the framework of infinite horizon control, this optimal movement duration would instead emerge from the controller properties without a priori calculation, as for Fitts’ law. Here, we expanded the infinite horizon approach used to capture Fitts’ law and determine whether it could be generalized to explain the tradeoff between reward and movement velocity.

A critical limitation of finite horizon models is their reliance on static conditions, assuming a fixed environment and movement duration for the determination of the control policy [3], [5], [16]. This contrasts with human reaching movements, which can rapidly be updated online in response to unexpected perturbations [18], [19]. To account for such flexibility, finite horizon controllers rely on computationally expensive ad hoc re-computations of their control policy online [18], [20]. In stark contrast, infinite horizon controllers offer a more natural solution through their inherent ability to optimize behavior over an infinite future, as shown by their ability to predict the increase of movement duration in presence of perturbations [6], [20], [21]. However, previous implementations fell short of capturing some specificity of human motor behavior, such as the temporal evolution of feedback responses [20], [22]. Here, we demonstrate that an infinite horizon control coupled with a nonlinear plant dynamic captures multiple aspects of the flexible behavior observed during reaching movements in dynamical environments, including the previously unaccounted temporal evolution of feedback responses.

The present work builds on existing infinite horizon optimal feedback control implementations [6], [7], [23], and study their generalizability across diverse aspects of human reaching movements. We demonstrated that this control framework not only effectively captures Fitts’ law [7] but also the tradeoff between movement duration and reward, previously explained by pre-specified movement duration [12]. Importantly, we reproduced previously published features of infinite horizon control applied to linear models, and extended their applications to nonlinear models, which were not previously studied. We found that it captured important aspects of motor behaviors, including the effects of rewards, motor decisions, and nonlinearities. This suggests a reconsideration of current evidence for finite movement time and proposes an alternative view regarding the origin of the modulation of movement duration in reaching movements as an emerging property of the selected control policy.

## Methods

### Problem formulation

We modeled the system dynamics using stochastic differential equations that capture how the state vector *x* evolves in presence of the motor commands *u*. This system dynamics is described by *dx* = *f*(*x, u*)*dt* + *Gdω*, where *f*(⋅) is a function capturing the limb dynamics and the action of the motor commands, *ω* is a realization of a standard Wiener process to express motor noise. The state vector is only observable through noisy sensory feedback *y* governed by *dy* = *g*(*x, u*)*dt* + *Ddξ*, where *g*(. ) is the observation function and *ξ* is another, independent realization of a Wiener process to express sensory noise.

### Nonlinear limb dynamics

We modeled the limb dynamics using a nonlinear dynamical system representing the planar movements of the arm as a system composed of two hinge joints corresponding to the elbow and shoulder joints. This model is referred to as the *two-joints model* and is simplified version of the models used in [24], [25]. The evolution of the elbow (*θ*_1_) and shoulder (*θ*_*2*_) angles, summarized by *Θ* = [*θ*_1_, *θ*_*2*_]^*T*^, is described by the Newton-Euler equations, which captures how the angular accelerations vary as a function of the inertia, the applied torques *T*, the Coriolis forces 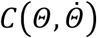, and damping forces. The system dynamics is described by: 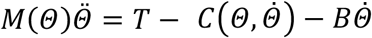, where *M*(*Θ*) is the inertia matrix and *B* is a viscous constant. We related the torques *T* to the motor commands through a first-order low-pass filter 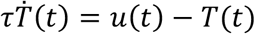, with a time constant of *τ* = 60*ms* to approximate the effect of nonlinear muscle dynamic [26]. The inertia and Coriolis matrices are expressed as follows

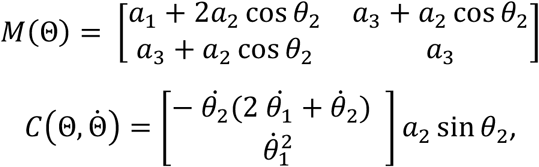

where *a*_1_, *a*_*2*_, and *a*_3_ are inertia terms defined based on each segment’s length, moment of arm and mass.

By defining the state vector as 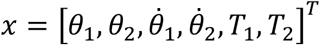, we can express this two-joint model as a nonlinear stochastic differential equation: *dx* = *f*(*x, u*)*dt* + *Gdω* . Importantly, this equation is nonlinear as both the inertia matrix and the Coriolis forces contain nonlinear dependencies on the angles and their derivatives. This system can be locally linearized using the Jacobian matrix evaluated at the current state *x* and motor command *u*, similar to [24], [25].

### Controllers

Most of the results presented in this study were obtained using an infinite horizon [7] formulation of optimal feedback control, a finite horizon [5] formulation of optimal feedback control was used for part of the results presented in the last figure.

### Infinite horizon optimal feedback control

We considered a system whose dynamics is described by a linear stochastic differential equation, *dx* = *Axdt* + *Budt* + *Yudγ* + *Gdω*, with the state observable through *dy* = *Cxdt* + *Ddξ*, where *dγ, dω*, and *d*ξ are Wiener processes. We searched the optimal motor commands *u*^∗^ that minimize a cost-functional reflecting both motor performance (i.e., the accuracy of reaching the target) and motor cost (i.e., the effort required) by adapting the controllers derived in [6]. By introducing a new variable 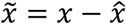, the stochastic differential equations for the state dynamics and observation can be coupled as 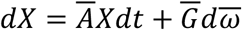, where 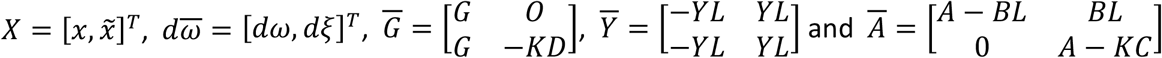.

The optimal states estimates are those that minimize the steady state estimation error 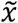, such that they minimize the cost-functional 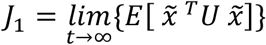, where *U* is a real-valued matrix. The optimal motor commands are those that minimize the cost-functional combining motor performance and motor cost 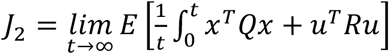, where *Q* and *R* are real-valued matrices. These two cost-functionals *J*_1_ and *J*_*2*_ are defined over an infinite time horizon. Based on the implementations developed in the previous work [7], [23], we define the cost-to-go matrix 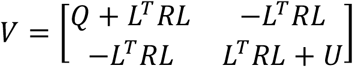 and the covariance matrix of the coupled state 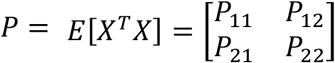. With these definitions, the combined functional *J* and the coupled stochastic differential equations are expressed respectively as 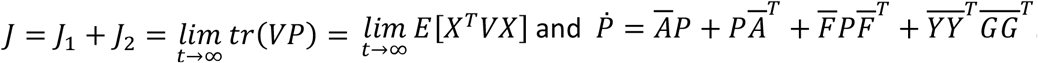.

Due to the presence of state-dependent noise, the controller and estimator gains cannot be solved independently. The control problem can be solved by minimizing the coupled cost-functional while satisfying the constraints imposed by the stochastic differential equations governing the state dynamics. This was achieved by introducing a Lagrange multiplier 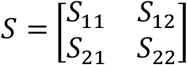. The solution of this optimization problem is given analytically by:

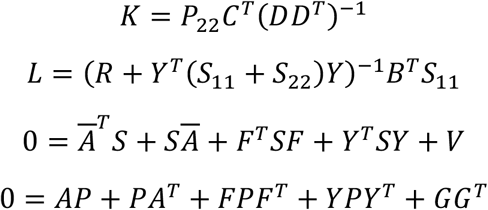

Since we solved the steady-state control problem, the feedback gains *L* and the Kalman gains *K* are time independent and remain constant throughout simulations. In practice, we obtained the optimal gain matrices by iterating the above set of equations until convergence. We initialized the iteration with the gain matrices from the finite horizon controller (evaluated at half of the movement duration), as the convergence was sensitive to the initial values of *K* and *L*.

### Finite horizon optimal feedback control

For the simulations involving the finite horizon controller, we considered the same dynamical system as for the infinite horizon formulation without signal dependent noise. We searched the optimal motor commands *u*^∗^ that minimize a cost-functional capturing both motor performance and motor cost over a finite time horizon; 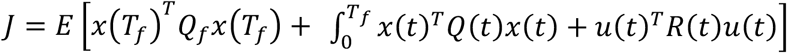, where *Q*_*f*_, *Q*(*t*), and *R*(*t*) are real-valued matrices representing the final motor performance, time-varying motor performance, and time-varying motor cost, respectively. Under the assumption that only the final motor performance term *Q*_*f*_is non-zero and that the motor cost *R* is constant and time independent, a closed-form expression for the time varying motor commands *u*^∗^ and states estimates 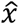 can be derived. These are given by 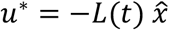 and 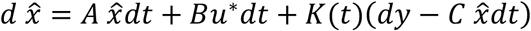, where *L* and *K* are the time-varying feedback and Kalman gain matrices.

To solve this control problem, we discretized the state dynamics to obtain the difference equation 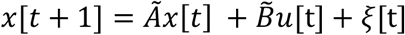, where *Ã* = *Adt* + *I* (with *I* as the identity matrix) and 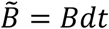, obtained using Euler method. The Wiener processes *dω* and *dξ* are discretize by the multivariate Gaussian noises *ξ*(*t*) ∼ *N*(0, *Σ*_*m*_) and *ω*(*t*) ∼ *N*(0, *Σ*_*s*_). We then derived the recurrence equations that define the optimal feedback and Kalman gains using Dynamic Programming [5]. In the absence of signal-dependent noise, the optimal controller and estimator can be resolved independently, and the corresponding gain matrices can be obtained recursively as follows,

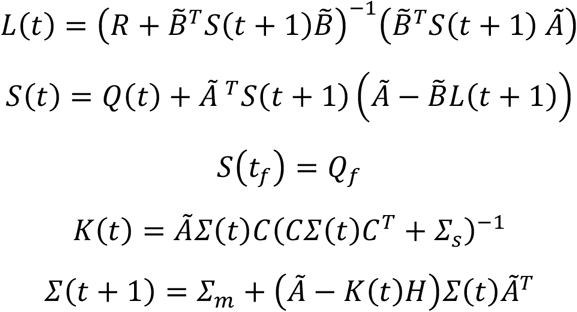

Importantly, the gains *L*(*t*) and *K*(*t*) derived with this finite horizon controller are time-dependent and thus vary throughout a single simulation.

### Simulating nonlinear dynamics with OFC

To apply the Infinite Horizon Optimal Feedback Control (IHOFC) to a nonlinear dynamical system, we need to linearize its dynamics. For finite horizon controllers, this problem was previously addressed by performing a local linearization around a nominal time-varying trajectory which was obtained by solving a deterministic version of the finite horizon control problem [24], [25]. The resulting controller operated within a tube around this nominal trajectory, with the assumption that the system remained in the vicinity of this nominal trajectory for the entirety of the simulation. This last assumption was not suitable for the movements we simulated here as it was very unlikely that the system remained in the vicinity of the nominal trajectory after being perturbed.

To address these challenges, we developed a variant of the method proposed by Todorov and Li [25] by implementing the state dependent Riccati equation method [27]. At each time step, we evaluated the Jacobian matrix of the system to linearize it around the current state; then solved the resulting linearized control problem for an infinite horizon, and finally executed one simulation step using the current control signal (Algorithm 1). We repeated this process until the end of the simulation time, fixed at 2s (200 timesteps) leaving enough time to the controller for stabilizing at the target. By continuously linearizing the system locally, the controller was able to adapt freely within the state space without being constrained in the vicinity of a fixed nominal trajectory.

Formally, the nonlinear state dynamics is captured by: *dx* = *f*(*x, u*)*dt* + *Gdω*, where *f*(*x, u*) captures the nonlinearities. The system dynamics can be linearized around the current state and motor command (*x*_0_, *u*_0_) by developing its first order Taylor expansion

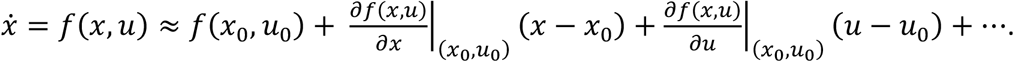

By introducing small deviations from the current operating state, *δx* = *x* − *x*_0_, and motor command, *δu* = *u* − *u*_0_, we can capture the local nonlinear behavior as

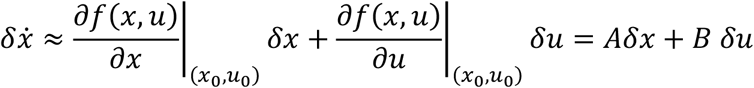

Since we did not linearize around a nominal trajectory obtained through open loop solution of the control problem, we do not have a nominal motor command *u*_0_, hence we considered *u*_0_ = 0 for our linearization. Note that this choice does not affect our linearized *A* and *B* matrices as the state dynamics of our system is linearly dependent on the motor commands.

The iterative algorithm we used for applying infinite horizon control to a nonlinear plant writes as follows:

#### Algorithm 1

**Iterative IHOFC control**

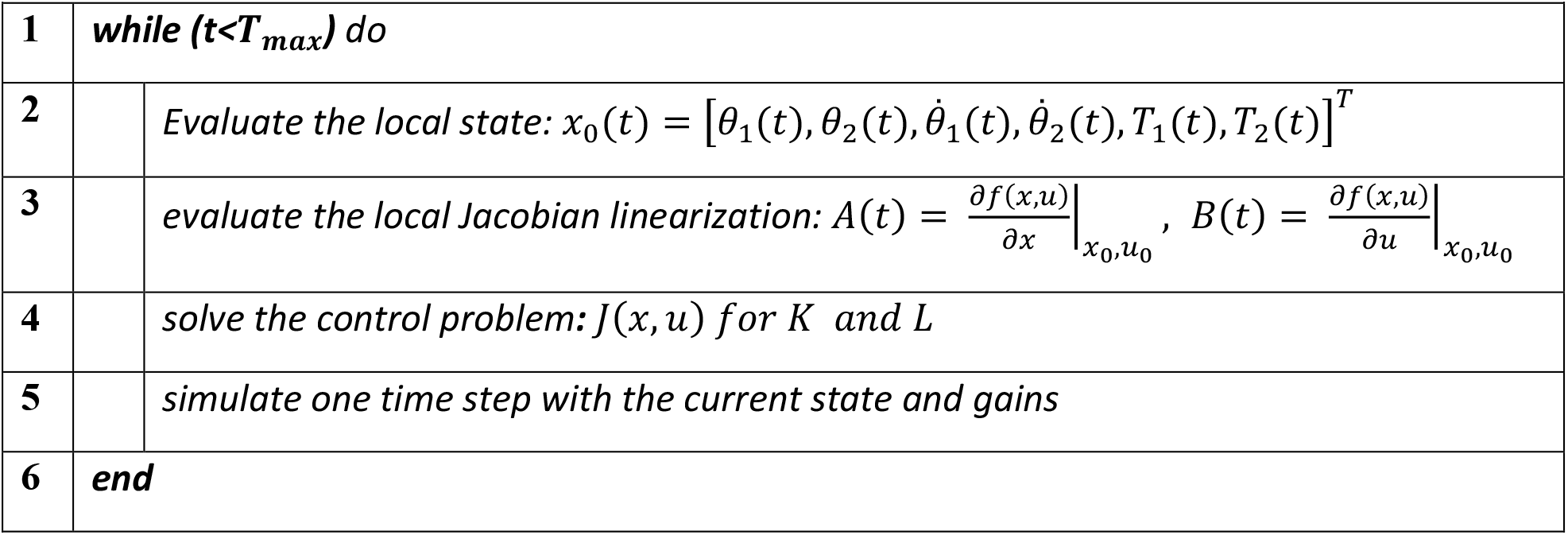

### Simulation varying reward

To investigate the controller’s behavior in the presence of reward, we incorporated a target reward into the cost-functional of the optimal problem. Experimentally, target reward influences participant’s tradeoff between effort and motor performance, leading them to engage in more effortful action when a larger reward is present [10], [28]. Similarly, previous simulation work suggested that reward acted by biasing the tradeoff between motor cost and motor performance [15], [29]. We implemented this influence by adding a diagonal matrix to the first entry of the matrix *V*, which corresponds to the controller cost-to-go, to simulate movements towards more rewarding targets. Crucially, this term was added to all the diagonal elements therefore differing from a modulation of the required accuracy only, which will target specific entries of that matrix instead [30]. This additional term increased the relative importance of motor performance compared to motor cost, resulting in a controller that is more prone to take effortful actions. The modified V matrix is given by:

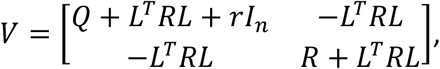

where *r* is the reward bias, which we varied within each experiment, and *I*_*n*_ is the identity matrix of dimension *n*.

### Determining movement duration

In all our simulated experiments, we used a consistent method to determine the movement duration for each condition. We defined movement duration as the time required for various variables – such as distance to the target, hand velocity, and the variability of hand position relative to the target (same as [7]) to fall below thresholds indicative of the required accuracy. The position and velocity conditions are straightforward and were met when 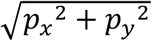 was less than the position threshold and 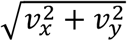 was less than the velocity threshold, both expressed in cartesian coordinates. For the condition related to the variability of hand position relative to the target, we used the first entry of the uncentered covariance matrix of the coupled state *P* = *E*[*XX*^*T*^], which corresponds to the uncentered covariance of the hand position. Movement duration is defined as the first time at which this uncentered covariance felt below the accuracy threshold for the first time.

## Results

We implemented an infinite-horizon optimal feedback controller (IHOFC) to determine whether the modulation of movement duration emerged from the control policies related to movement execution. Our approach, based on previous work [6], [7] was adapted to test various experimental conditions. First, we investigated whether the ability of the IHOFC to reflect the modulation of movement duration with target distance and accuracy (Fitts’ law, [2]) holds when considering nonlinear state dynamics and different criteria for movement duration. Second, we investigated whether IHOFC captures the impact of reward on movement vigor and decision-making [1], [9], [10], [11]. Finally, we studied to what extent considering a nonlinear model of the limb dynamics helped capture human behavior in presence of perturbations [22].

### Modulation of movement duration with task parameters

We began by investigating the IHOFC’s ability to capture how movement duration was modulated by target distance and accuracy constraints, in line with Fitts’ law. More specifically, we sought to determine whether the properties of the IHOFC controller [7], [8] hold when considering nonlinearity and experimentally realistic stopping criteria. For this purpose, we simulated reaching movements towards targets located at different distances from a starting position (Figure 1A). We numerically defined movement duration as the first crossing of thresholds on different movement variables: the expected square of the hand position as a measure of uncentered variability (Figure 1E), distance to target (Figure 1F), and hand velocity (Figure 1G). We varied these thresholds to simulate variations in accuracy levels.

**Figure 1.**
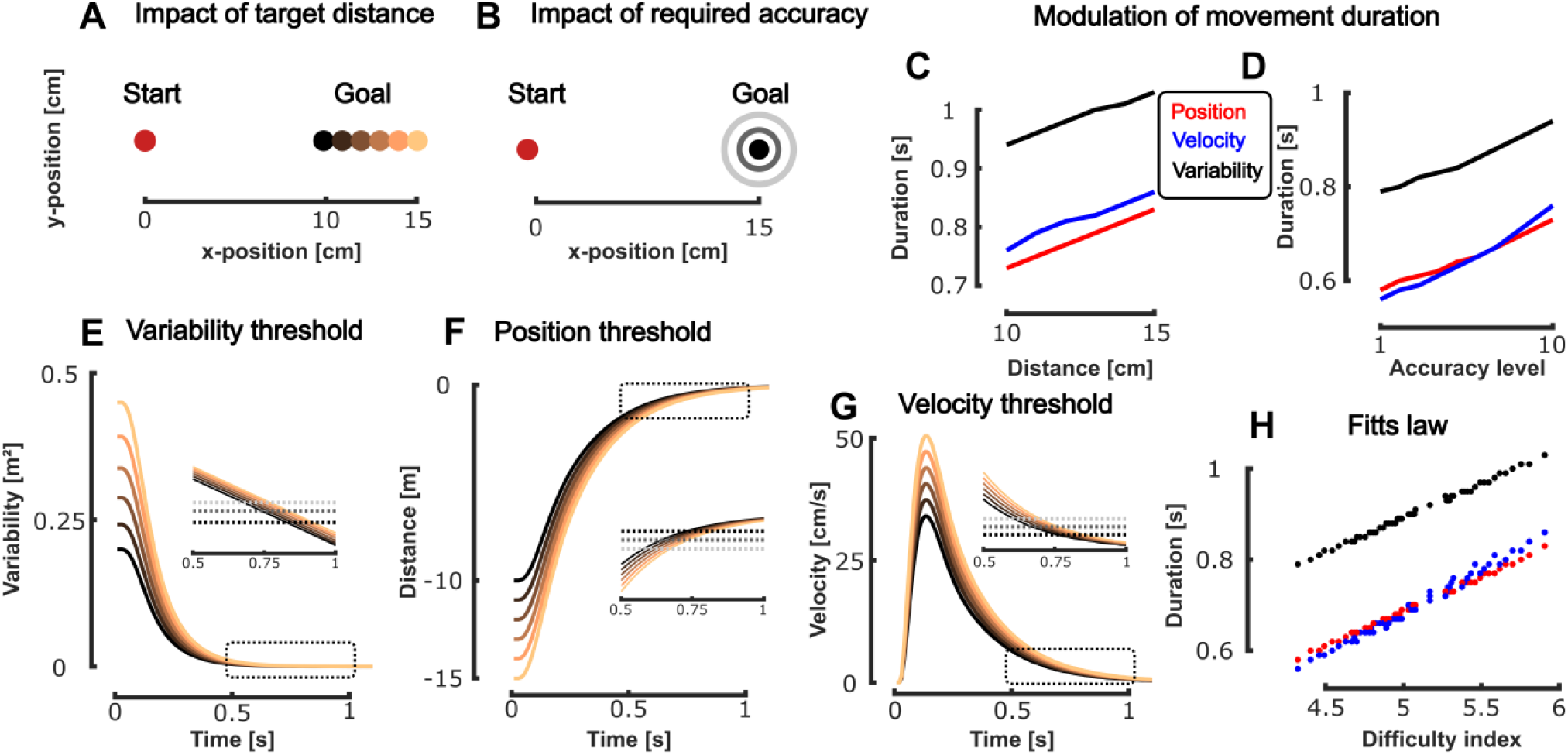
Modulation of movement duration with task parameters for a nonlinear dynamical system – Modulation of target distance (**A**) and required accuracy (**B**), same colormaps as in panels **E-G. C** Modulation of the movement duration as a function of the distance to the goal target for the position (red), velocity (blue) and variability (black) accuracy-related thresholds. **D** Modulation of the movement duration as a function of the required accuracy for the position (red), velocity (blue), and variability (black) accuracy-related thresholds. Evolution of the uncentered position variance (**E**) (with respect to the goal target), distance (**F**), and velocity (**G**) as a function of time for the different target distances (color coded). Movement duration is defined as the crossing point between the different metrics and the threshold. **H** Illustration of the relationship between the movement duration and the difficulty index for the position (red), velocity(blue) and variability (black) accuracy-related thresholds.

The nonlinear IHOFC model predicted an increase in movement duration for more distant targets with all three thresholding variables (distance, velocity, and uncentered variability in red, blue, and black in Figure 1C, see Methods), consistent with human behavior and prior simulations [7], [8]. Similarly, IHOFC predicted an increase in movement duration with higher accuracy requirements (Figure 1D), aligning with past experimental and modeling work. The absolute value of the movement duration was influenced by the thresholding method but its relationship with target distance and accuracy requirements remained similar across all three methods (Figure 1E-F). The combination of the scaling of the trajectories with the superposition principle and the use of a fixed threshold to determine the end of movement produced this apparent modulation of movement duration.

Movement duration was simultaneously modulated by the distance to the target and the required level of accuracy as captured by the speed-accuracy trade-off defined by Fitts’ law. This trade-off was captured by a linear relationship between movement duration and the logarithm of a difficulty index defined as the ratio between the distance and required level of accuracy. We defined this difficulty index as in previous work [7], [15] and simulated reaching movements while varying both the target distance and the required accuracy (captured by the cost matrices). We observed that the movement durations associated with each pair of parameters were linearly dependent on the difficulty index for all three thresholding methods (Figure 1H), as predicted by the speed-accuracy trade-off. We observed the same results for a linear dynamical system (Figure S 1). These results showed the ability of the IHOFC model to capture a fundamental aspect of movement duration during reaching (Fitts’ law), even when considering a nonlinear system dynamic, and revealed that the absolute value of movement duration may result from a by-product of numerical criteria defined during offline processing.

### Modulation of movement duration with reward

We then investigated whether the IHFOFC model could capture how increasing target reward yielded to a decrease in movement duration [10], [31] and a concomitant increase in feedback responses [32], both reflecting a modulation of movement “vigor” .We varied the target reward by increasing the square penalty associated with rewarding targets (see Methods), and simulated unperturbed reaches across a range of reward values. The IHOFC reproduced the fact that target reward influenced the hand trajectories (Figure 2A) and reduced the movement duration in presence of higher reward values (Figure 2B). The motor cost associated with movements, defined as the sum of squared commanded torques, increased for the shorter movements in high-reward conditions (Figure 2C), consistent with the observation that humans and animals are willing to perform costlier actions when their motor costs are counterbalanced by more rewarding states [12]. This increase was observed for both the elbow and the shoulder joints (Figure 2D-E).

**Figure 2.**
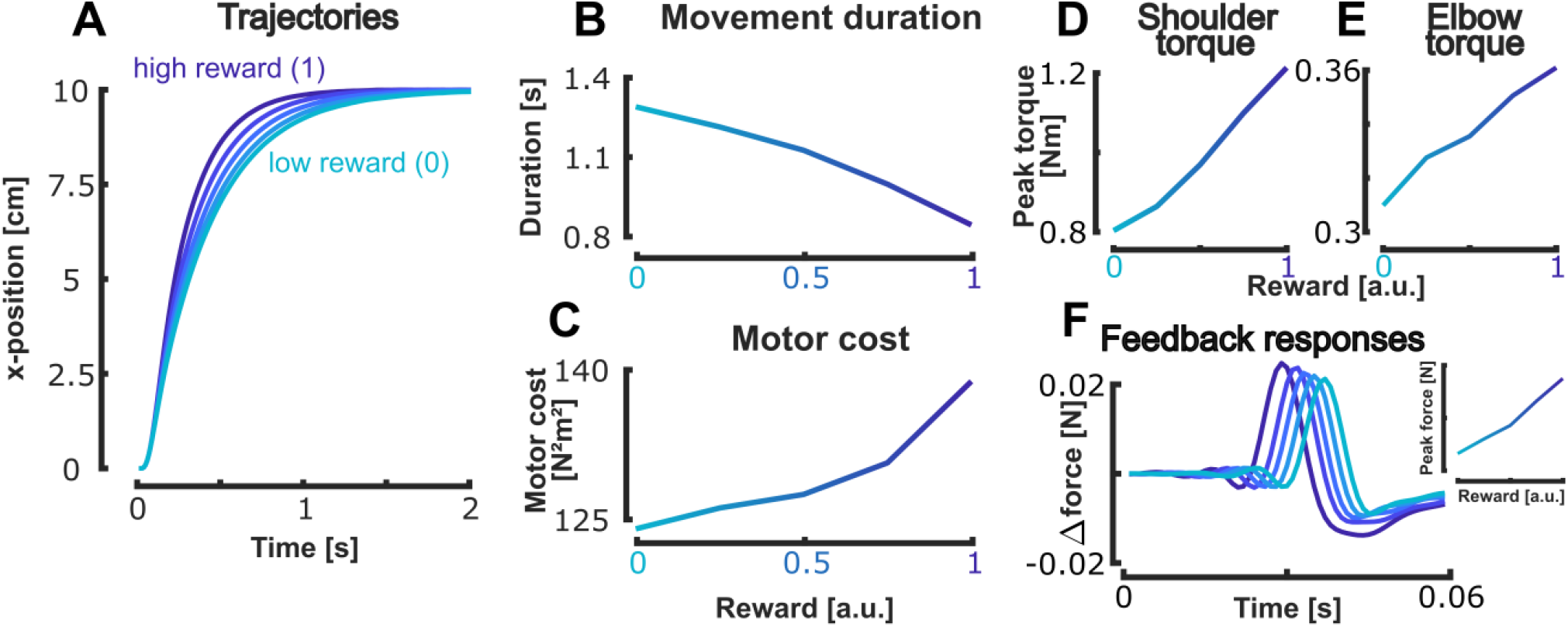
Modulation of movement duration with target reward - **A** Evolution of the distance to the target as a function of time for the different reward condition. Darker and lighter blue traces respectively capture larger and smaller reward, respectively. Modulation of movement duration (**B**) and motor cost (**C**) with the target reward. The leftmost and rightmost dots respectively represent the low and high reward conditions. Peak shoulder (**D**) and elbow (**E**) torques as a function of reward for the unperturbed conditions. **F** Modulation of the feedback responses to target jump as a function of the target reward (color coded). The insert captures the peak responses for each reward condition.

We then assessed the model’s response to perturbations by simulating changes in target location during movement. For each reward condition, we moved the target location by 3cm in the positive y-direction once the hand had moved for half of the distance to the target. We observed that the amplitude of feedback responses, expressed as the increment of force in presence of perturbation, increased with increasing reward value (Figure 2F). Combined, these results reproduce a range of empirical experimental findings from human reaching studies [31], [32], [33], indicating that the IHOFC model also captured the effects of reward on movement vigor for both goal-directed movements and feedback responses [14]. We observed similar results with a linear dynamical system (Figure S 2).

We next explored the IHOFC’s ability to capture how reward influenced decision-making when participants could choose which target to reach amongst multiple targets [31], [34]. To decide which target to reach, humans balance the cost of effort and the potential reward outcome of each action, i.e. a reward-motor cost trade-off [35]. Here, we simulated a task where participants had to reach for any of two targets aligned vertically (Figure 3A). The vertical position of the top target was fixed at 3cm from the starting location, while the bottom target position was varied from the symmetric location of the top target to a maximum vertical distance of 7.5cm. The reward of the bottom target was set to 0 and kept constant across conditions, while we varied the reward of the top target. For each pair of target distance and reward, the reached target was selected as the one associated with the lowest cost-to-go value, as done in previous work [29]. The IHOFC predicted that the selected target varied with both the target locations and rewards (Figure 3B), capturing respectively the influence of motor cost and movement outcome. If the top target was associated with a small reward (leftmost column in Figure 3B), it was only reached when the bottom target was very distant. In the case illustrated in Figure 3C, the bottom target (blue) was selected as long as it was within 5cm of the vertical position of the starting location, the top target (yellow) was selected if the bottom target was at more than 5cm of this location (the black arrow in Figure 3B illustrates this transition). Moreover, our model predicted that movement duration was still modulated with the target reward (Figure 3D, illustrated for the conditions highlighted by the full line red box in Figure 3B) and the target distance (Figure 3E, illustrated for the conditions highlighted by the dashed line red box in Figure 3B). Note that the modulation of movement duration in the IHOFC simulations was not related to the time elapsed in the decision process, and purely related to how the movement was executed once a decision has been made. The results presented in Figure 3 were obtained for a linear plant dynamic, similar to the one used in [29], because of the amount of computation required to generate the results with a nonlinear system, we however obtained similar results in that later condition when tested on a smaller version of the task (Figure S 3).

**Figure 3.**
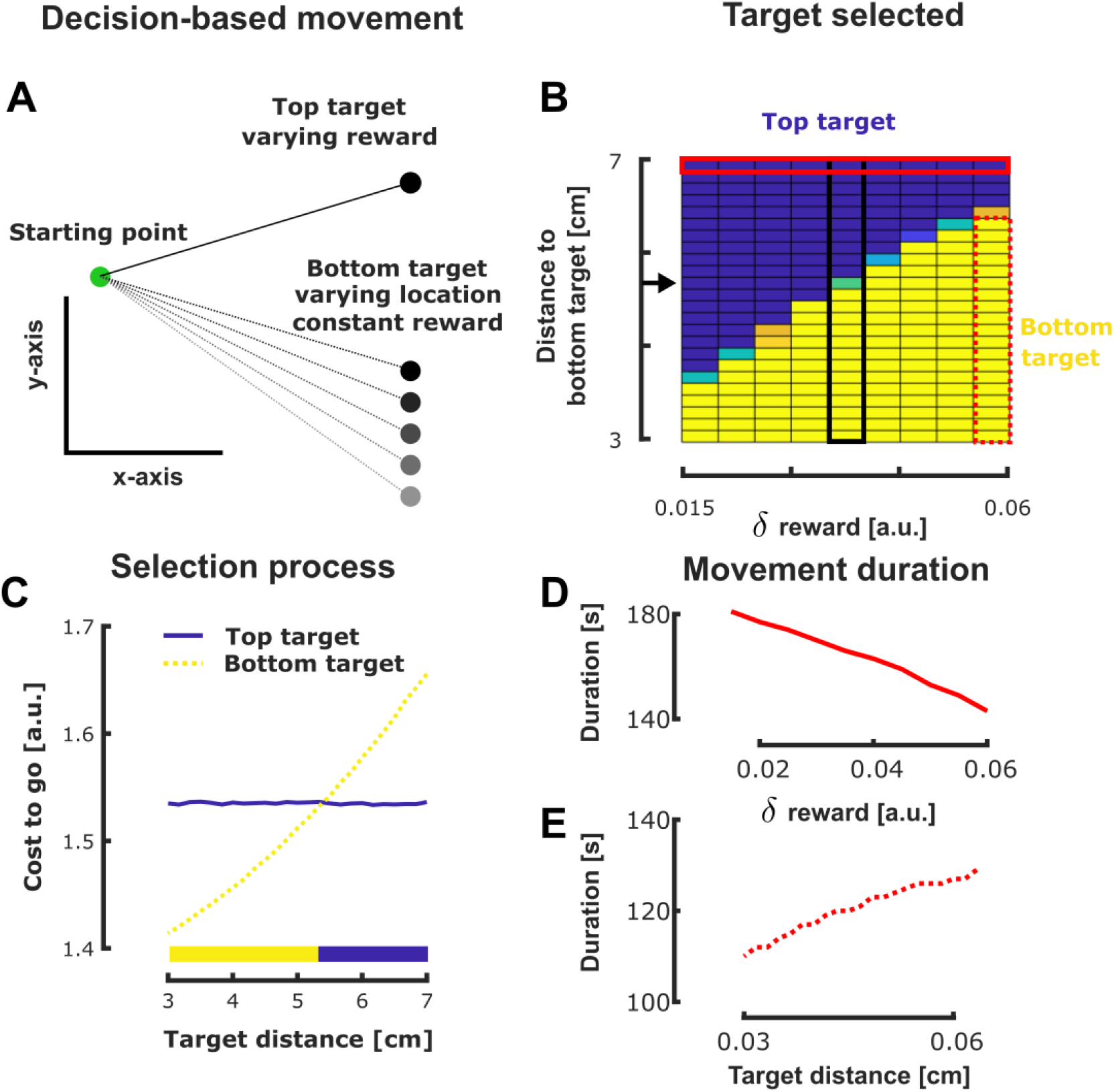
Illustration of the decision-making processes in presence of reward and varying target location. **A** Schematic representation of the simulated experiment. We simulated reaching movements from the starting point (green) to two different targets (top and bottom). We varied the position of the bottom target and the reward associated with the top target across the trials. **B** Heatmap of the target reached for each pair of top target reward and bottom target distance (blue and yellow capture movements towards the top and bottom targets respectively). **C** Illustration of the cost-to-go (in arbitrary units) based decision process for the conditions highlighted by the black box in panel **B**. Blue and yellow lines capture the cost-to-go as a function of the bottom target location for the top and bottom targets respectively, for a fixed reward difference. The colored rectangles indicate the selected target. **D** Modulation of movement duration as a function of the top target reward for the conditions highlighted by the red full line box in panel **B. E** Modulation of movement duration as a function **of the bottom target location for the conditions highlighted by the red dashed line box in panel B**.

### Benefits of a nonlinear plant model

We finally explored the benefits of using a nonlinear system dynamic with an IHOFC. Linear IHOFC captures the modulations of movement duration with a wide range of task parameters but they have the major caveat that their feedback gains are constant, as they are obtained by solving the steady-state control problem [7]. In contrast, nonlinear IHOFC has feedback gains that depends on the current state (see Methods). We therefore wondered whether it would be enough to capture the modulation of movement duration with motor cost for equidistant targets [36], [37], [38] and the temporal evolution of feedback responses during movements [22], that that linear IHOFC models failed to capture [20].

We first simulated center out reaching movements to investigate whether the IHOFC model could capture the trade-off between movement duration and the effort cost associated with that movement. We hypothesized that the IHOFC would take more time to reach the target in more challenging conditions, similarly to what we observed when increasing the level of required accuracy. To verify this hypothesis, we compared the movement duration to the different targets in the center-out reaching task (Figure 4A) and computed the associated motor cost, expressed as the sum of squared actuated torques. As we hypothesized, we observed that the targets associated with larger motor costs were reached the slowest (Figure 4B).

**Figure 4.**
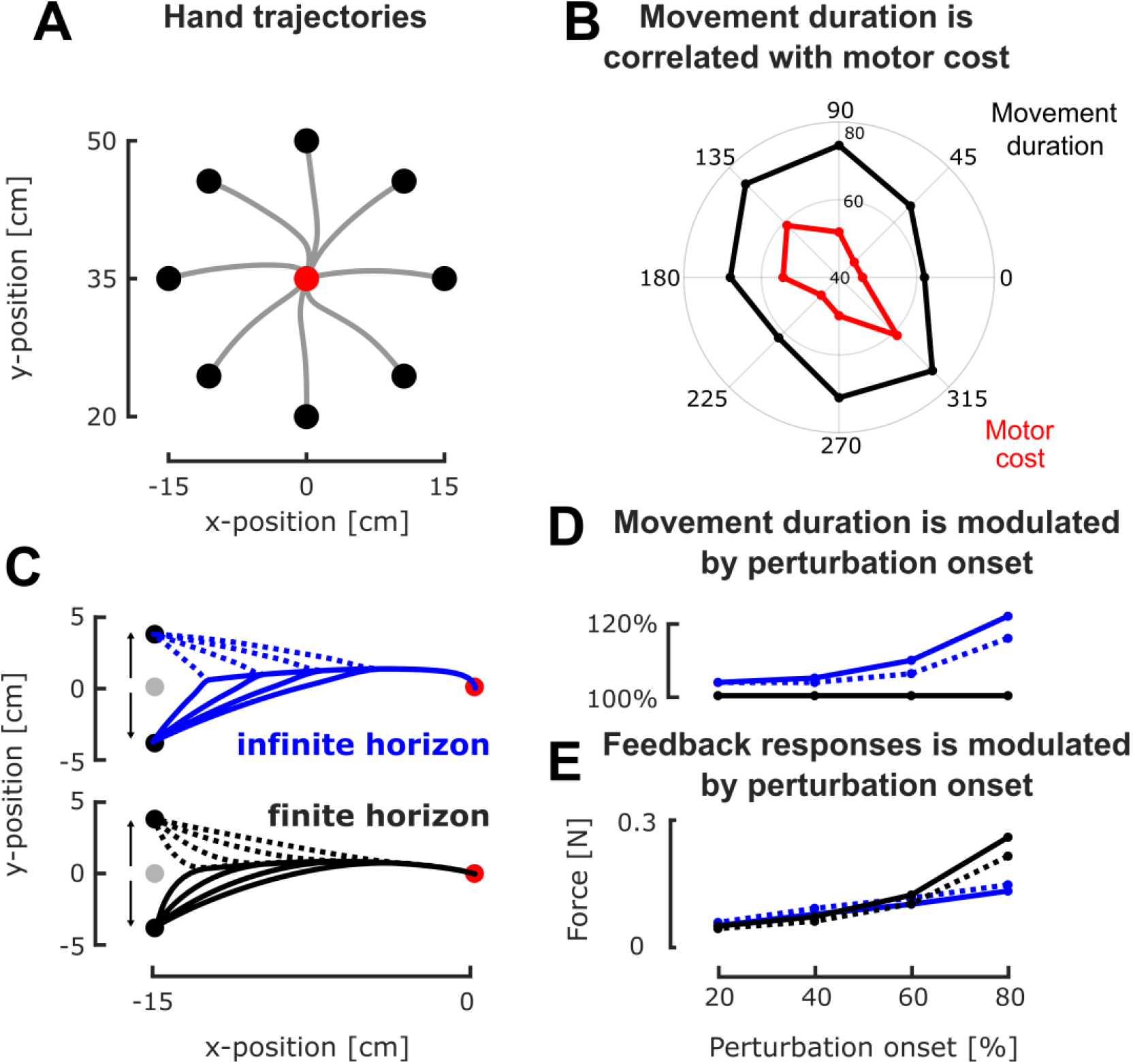
Modulation of movement duration for a nonlinear system - **A** Simulated hand trajectories for center-out reaching movements between the starting position (red) and the different goal targets, using the IHOFC controller. **B** Correlation of the movement duration (black, radial axis) with the total motor cost (red, radial axis) for the different targets in the center out task. Targets associated with large motor costs were reached slower than those associated with low motor costs. **C** Hand trajectories for reaching to the leftmost target of the center out reaching task (cyan target) in presence of target jumps for the infinite (top) and finite (bottom) horizon nonlinear controller. Full and dashed lines capture counterclockwise and clockwise perturbations respectively. **D** Modulation of movement duration - expressed relative to the unperturbed movement duration - as a function of perturbation onset for the infinite (blue lines) and finite (black lines) horizon controllers. **E** Modulation of the feedback responses to the target jump as a function of the perturbation onset for the infinite (blue lines) and finite (black lines) horizon controllers. Full and dashed lines respectively represent the counterclockwise and clockwise perturbations.

We then simulated reaching movements towards the same set of targets in presence of target jumps that could occur at 20, 40, 60, and 80% of the unperturbed movement duration (Figure 4A). The goal target was always located 15cm from the starting position and could jump 4.2 cm orthogonally to the main movement direction, either in clockwise (dashed lines in Figure 4C) or in counterclockwise (full lines in Figure 4C) direction. Our implementation of nonlinear IHOFC predicted an increase of movement duration with the occurrence of perturbations with larger increases for the later perturbations (Figure 4D, blue lines whereas a finite horizon optimal feedback controller (FHOFC) predicted constant movement durations (Figure 4D, black lines). Surprisingly, the feedback responses (measured as the increment of force orthogonal to movement direction) varied with the onset time of target jump with both the IHOFC (Figure 4E, blue lines) and FHOFC (Figure 4E, black lines). These results suggest that the IHOFC model can qualitatively capture the time-varying modulations of responses forces during movement if the controlled dynamical system is nonlinear. Thus, time-varying control responses may not be uniquely due to the presence of a time horizon, and it is important to consider both sources (nonlinearities and finiteness of the time horizon) when interpreting the time course of control responses to perturbations occurring at distinct moments.

## Discussion

In this work, we illustrated the properties of infinite horizon optimal feedback control as candidate theoretical framework to study human reaching movements. This framework captures different tradeoffs between movement duration and task parameters (Fitts’ law, reward-velocity) as well as the modulation of feedback responses to perturbations, without requiring a priori specification of movement duration that is necessary in finite horizon models. Together, our results broaden the explanatory scope of infinite horizon control and provide important insights to refine the ongoing debate between finite and infinite horizon models in movement neuroscience.

We were able to capture tradeoffs between movement duration and task parameters thanks to a property of the infinite horizon control framework: movement duration emerged directly from the control policy. For instance, while previous work using this framework captured Fitts’ law [7], [8], we reproduced these results and even extended them to the case of nonlinear plant dynamics based on relevant thresholds, such as hand position or velocity. We also demonstrated that this framework captured the tradeoff between movement vigor and reward, where movements associated with larger rewards were faster and exhibited more vigorous feedback responses [14]. In our simulations, the same property emerged directly without requiring to consider a temporal discounting of reward to define an optimal movement time [12], [15] . Furthermore, our model also captured the tradeoff between biomechanical cost and movement duration, whereby movements were slower when associated with a larger biomechanical cost [36], [37], [38]. While previous approaches, such as models based on expected utility, captured several of these tradeoffs [39], our work demonstrated their emergence within a unified optimal feedback control framework [5].

Infinite horizon controllers are also well suited to capture the flexibility of human reaching behaviors. For instance, our work highlights the infinite horizon controller’s inherent ability to explain human reaching behavior in response to perturbations. Unlike finite horizon control, which explicitly relies on a fixed movement duration determined for a static environment and requires ad hoc recomputations for mid-movements changes in movement duration, infinite horizon control naturally adjusts its movement duration during movement. Importantly, while previous modeling work using infinite horizon controllers captured the modulation of movement duration with perturbation [6], [20], [21], they did not explain why the feedback responses were modulated by the onset of perturbations [22], while finite horizon control models did [20]. Here, we captured this feature by using a nonlinear plant dynamic. Moreover, our work also captures how human behave when facing multiple targets [34], [40], [41]. Their decision occurs through a weighting of the reward outcome and motor effort [35] of each option. In simulations, this can be achieved by computing the cost-to-go associated with each option and selecting the control policy associated which whichever option has the lowest cost-to-go [29]. Here, we demonstrated that this methodology also works with infinite horizon controller and does not alter the modulation of movement duration with reward or distance. Together, our developments, capturing flexible aspects of human motor control, constitute a strong argument for infinite horizon control as a candidate theoretical framework for understanding human reaching movements.

Our main contribution is to propose a conceptual shift: movement duration may not be a pre-computed variable, but instead an emergent property of the controller’s specification. Previous interpretations of human behavior were rooted in theoretical frameworks assuming that movement duration is predefined and recomputed when necessary [18], [20], [29], while the infinite horizon controller presented in this work as well as in previous ones [6], [7] offers a less computationally demanding viewpoint. This contrast is nicely illustrated by the modulation of reaching behavior with reward. It is experimentally well established that reward increases movement vigor [10], [14], modulates feedback responses [31], [32], [33], and influences decision-making [31], [34], [40]. While these effects can be accounted for by combining a temporal discounting of reward and the cost-of-time [12], [13], [14], [15], our approach achieves these effects without such an explicit assumption. The underlying mechanism is different: since the optimal control gains scale with the cost parameters, increasing the cost of missing a target result in an increase in feedback gains, which modulates movement vigor. While we do not reject the possibility that a cost-of-time, sensitive to rewards, participates in the determination of movement time, we show that it may not be the sole mechanism by which rewards can modulate movement time.

While our infinite horizon model captured a wide range of phenomena, it is important to acknowledge certain limitations related to our current approach and its scope. One key limitation is the inability of the model to capture the target undershoot observed when reaching for a moving target [18]. This may be a signature of finite horizon controllers, which prioritize the velocity and effort penalties over the position term near movement end. Furthermore, our current infinite horizon formulation struggles with tasks requiring sequences of distinct movements to multiple targets [42], [43], [44] since it considers all subsequent targets and would therefore not reflect the limited planning horizon observed in humans. Methodologically, our implementation was restricted to target jump perturbations as numerical instabilities prevented us from studying mechanical perturbations, which future studies could expand. Finally, our methodological choice for the linearization relies on the local approximation of the known state dynamics which limits the quantitative interpretability of our results. The goal of that linearization was to demonstrate the robustness of the properties of the infinite horizon controller in presence of nonlinear state dynamics, and not to make assumptions about how the problem is solved in the brain. Future work could further investigate the nonlinear properties of those controllers by considering more advanced linearization methods or control models capable of directly control nonlinear systems. Specifically, our finding that the biomechanical cost associated with a target increased the movement duration (Figure 4B) offers a testable hypothesis that could be investigated in experimental work.

To conclude, we hypothesize that, in practice, humans likely leverage a wide range of control horizons depending on the context of the task: Finite horizons are likely suitable for familiar, automated movements, or in cases where an explicit timing constraint is imposed by the task, whereas infinite horizon is adequate in contexts where there is no explicit timing requirement (e.g., postural control, tracking etc.). Intermediate cases may certainly exist, where intermediate or long horizons appear similar to infinite horizons. While different tasks may be naturally associated with finite and infinite horizons, the originality of our developments was to show that the infinite horizon control scheme made non-trivial predictions about apparent changes in movement time even in the simple case of short, point-to-point reaches. This clearly highlights potential pitfalls in the intuitive distinction between those frameworks, and suggests that the assessment of controller horizon, or decisions about movement time, requires careful modelling work. The possibility that multiple control strategies might be available opens exciting future research directions. Towards designing experiments that could reveal whether and when humans use each control policy, finite, or infinite horizon, it will be important to thoroughly document the properties of both approaches to guide future investigations.

## Acknowledgments

A.D.C. was supported by a WBI Mitacs-Globalink grant from the Belgian government for this work and is currently supported by a postdoctoral fellowship from the K. Lisa Yang Integrative Computational Neuroscience (ICoN) Center at MIT. H.T.K was supported by the Fonds de la Recherche Scientifique (F.R.S.-FNRS) Chargé de recherche Grant CR 252 (FC 043127). J.A.P was supported by the Canada Research Chairs Program.

## Supplementary material

### S1. Fitts’ law with linear dynamics

**Figure S1.**
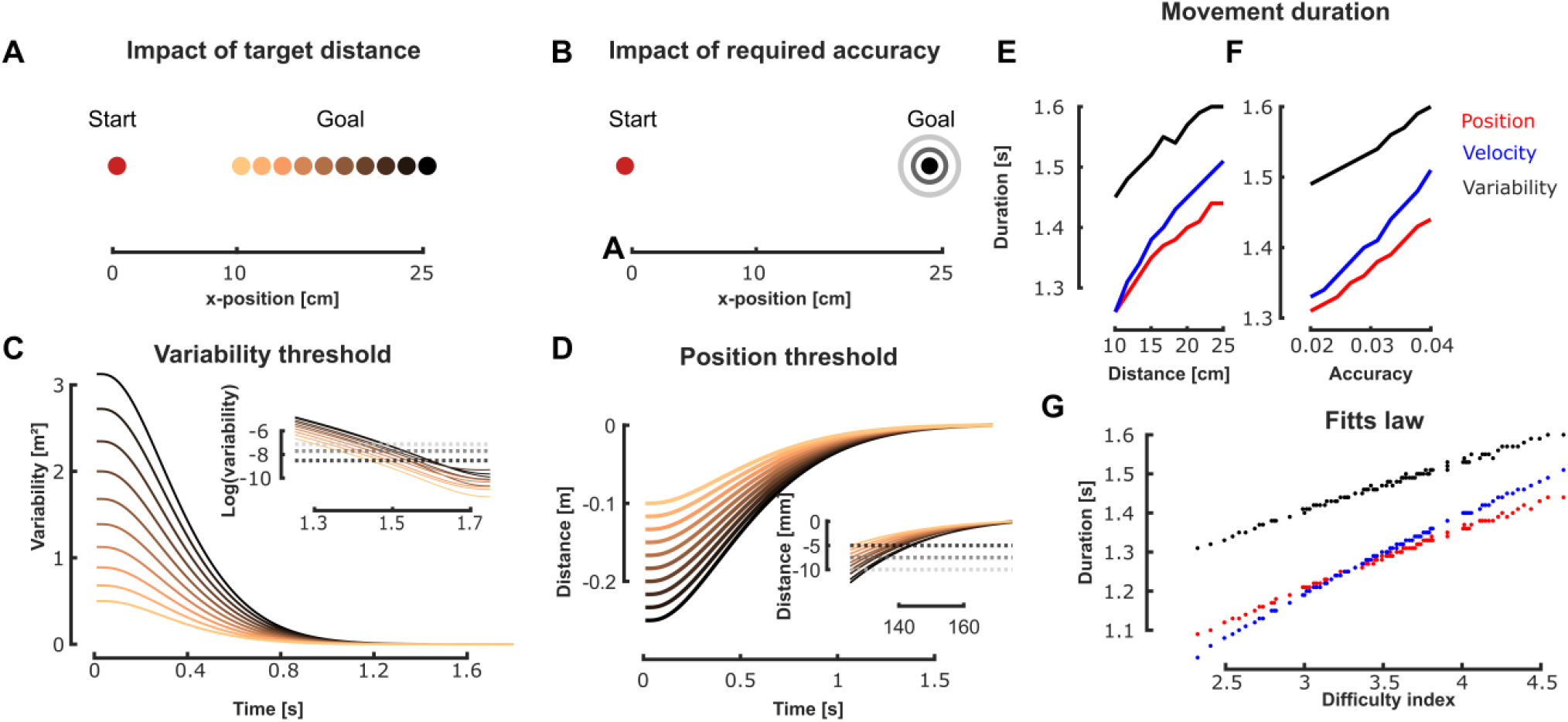
Modulation of movement duration with task parameters - **A** Schematic representation of the modulation of the target distance. The red dot represents the starting position, and the different colored circles represent the different goal targets. **B** Schematic representation of the modulation of the required accuracy. The concentric circles located around the black goal target represent decreasing requirement of accuracy captured in the cost matrices. **C** Evolution of the uncentered position variance (with respect to the goal target) as a function of time for the different target distances (color coded). The insert is an enlargement that represents the accuracy-related variability thresholds (grayscale) for three different accuracy levels. Movement duration is defined as the crossing point between the variability and the threshold. **D** Evolution of the distance to the target as a function of time for different target distances (color coded). The insert is an enlargement that represent the accuracy-related position thresholds (grayscale) for three different accuracy levels. **E** Modulation of the movement duration as a function of the distance to the goal target for the position (red), velocity (blue) and variability (black) accuracy-related thresholds. **F** Modulation of the movement duration as a function of the required accuracy for the position (red), velocity (blue), and variability (black) accuracy-related thresholds. **F** Illustration of the relationship between the movement duration and the difficulty index for the position (red), velocity(blue) and variability (black) accuracy-related thresholds.

### S2. Impact of reward with linear dynamics

**Figure S2.**
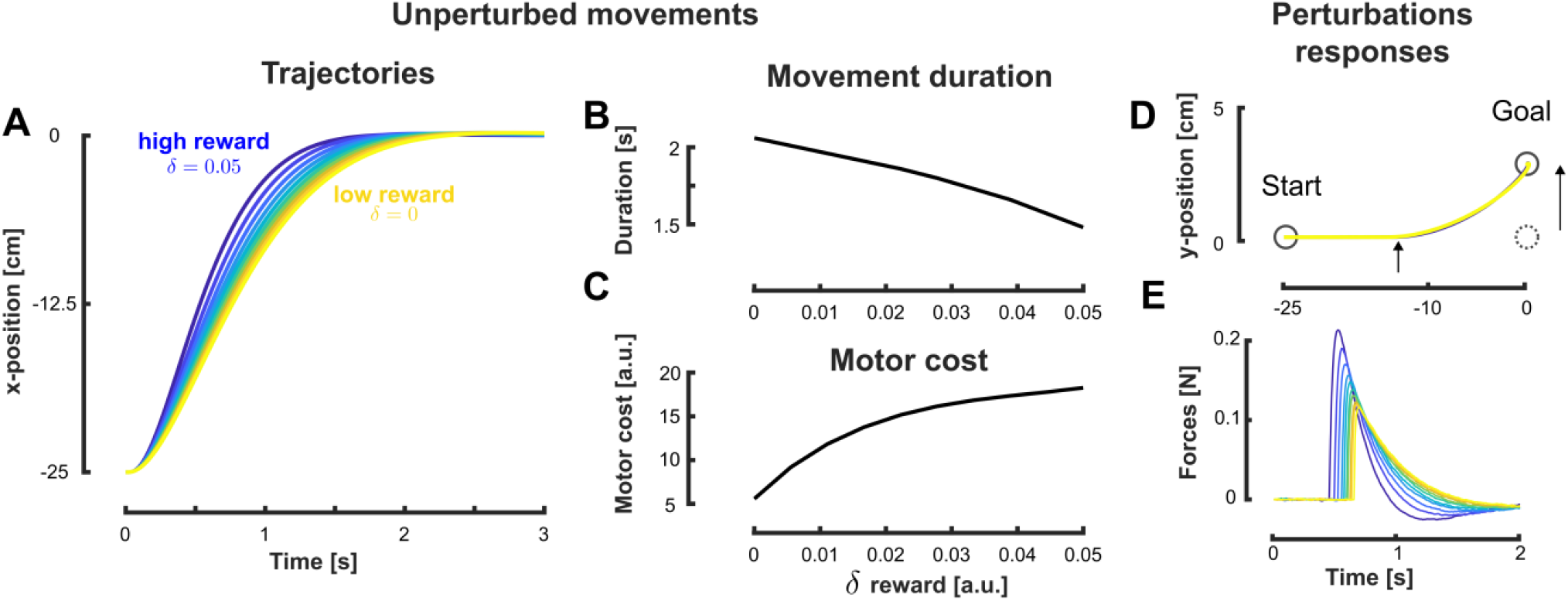
Modulation of movement duration with target reward - **A** Evolution of the distance to the target as a function of time for the different reward condition. Blue and yellow traces respectively capture larger and smaller reward. **B** Modulation of movement duration with the target reward. The leftmost and rightmost dots respectively represent the low and high reward conditions. **C** Modulation of the motor cost with the target reward. The leftmost and rightmost dots respectively represent the low and high reward. **D** Average simulated hand trajectories for the larger (blue) and smaller (yellow) reward values. The target changed location during movement, the small arrow illustrates the onset of target jump. **E** Modulation of the maximal feedback responses with the target reward. The leftmost and rightmost dots respectively represent the low and high reward.

### S3. Decision making in presence of nonlinear dynamics

**Figure S3.**
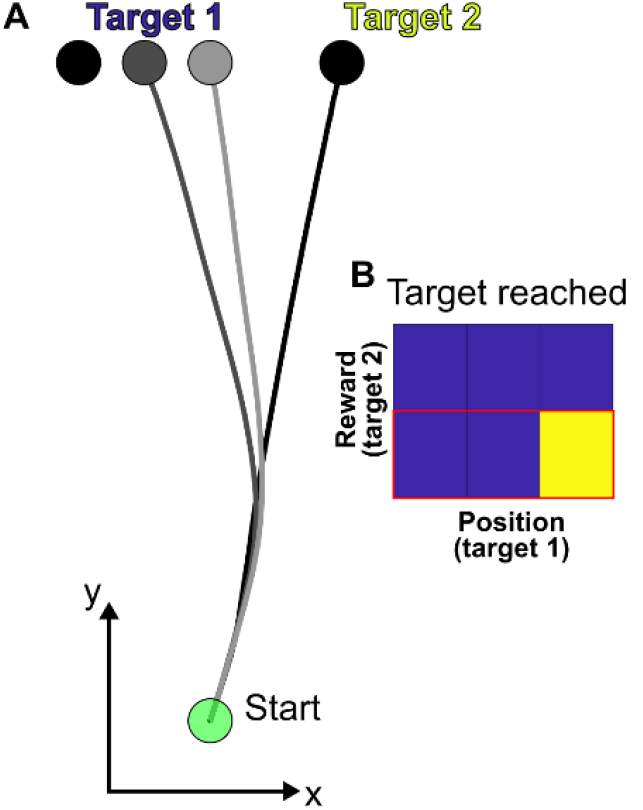
Decision-making in for a nonlinear plant dynamic. (**A**) Sampled trajectories for three different conditions where the position of target 1 was varied (color coded) while target 2 remained at the same location. (**B**) Target reached for different combinations of target 1 location and target 2 reward. Blue and yellow boxes respectively represent conditions for which targets 1 and 2 were reached. The row highlighted in red corresponds to the sampled trajectories of panel **A**.

